# Cardiac CapZ regulation during acute exercise

**DOI:** 10.1101/2020.03.03.975185

**Authors:** Andrew Laskary, Logan K. Townsend, David Wright, W. Glen Pyle

## Abstract

**Purpose:** Exercise requires a rapid cardiac response to maintain cardiovascular function. CapZ is a critical stress-response protein in cardiac myocytes. While its role in the pathological stress response has been explored, its part in the physiological response to exercise is unknown. This study examined CapZ regulation during acute exercise and sought to determine its importance in the cardiac response to exercise.

**Methods:** Wildtype or cardiac CapZ-deficient transgenic mice were subjected to 20 min of swimming, or exhaustive exercise protocols. Time to exhaustion was a measurement of exercise capacity. Following submaximal exercise, cardiac myofilaments were isolated and probed for CapZ and key regulatory proteins. Myofilament function was assessed using an actomyosin MgATPase assay and total protein phosphorylation quantified with ProQ Diamond staining. Myofilament regulatory proteins following submaximal exercise were quantified by immunoblotting.

**Results:** Total myofilament CapZ was unaffected by exercise but increased total CapZIP and decreased phosphorylated CapZIP indicated weakened CapZ-actin interaction. CapZ-deficient transgenic myofilaments lacked changes in CapZIP but BAG3 increased 10%. Time to exhaustion was lower in CapZ-deficient mice in both swimming and running protocols. Actomyosin MgATPase activity was maintained in wildtype mice and impaired with CapZ deficiency. Exercise increased the phosphorylation of several myofilament proteins in wildtype mice but not transgenic animals. Exercise-dependent Increases in myofilament PKC-α and -ε were mitigated in CapZ-deficient mice. Tcap levels decreased 39 ± 8% in CapZ-deficient myofilaments with exercise and leiomodin 2 increased 78 ± 28% in wildtype myofilaments.

**Conclusions:** Cardiac CapZ is a critical player in the physiological response to exercise. CapZ-actin binding is rapidly altered with exercise. Decreased cardiac CapZ limits exercise capacity, impairs myofilament regulation, and leads to a less stable contractile apparatus.

## Introduction

The heart is a highly dynamic organ, responding rapidly to a range of environmental stimuli in order to match function with demand. For example, exercise increases heart rate and myocardial contractility within seconds of onset (1). The period of time associated with the physical activity was described by Fisher as the ‘acute’ phase, and is characterized by a series of molecular changes that drive adaptations which have profound short- and long-term implications on cardiac structure and function (2). Despite the acknowledged ability of the heart to respond quickly to exercise and the long-term impact the acute molecular alterations have, there is limited understanding of the intracellular events that underlie the functional transformation of the myocardium during aerobic exercise (3).

Chronic exercise regimens dramatically alter the architecture of the heart in large part through cellular hypertrophy. Much of the hypertrophic response involves alterations in gene expression that ultimately lead to both quantitative and qualitative proteomic changes. Although these structural alterations are chronic, taking weeks to reach equilibrium, there is evidence that the molecular structure of cardiac myocytes is affected much more rapidly. For example, physical stress applied to both skeletal (4) and cardiac (5) muscle rapidly alters actin filament capping by the Z-disc protein CapZ, leading to actin filament remodelling. We have shown that relatively small alterations in cardiac CapZ levels significantly impacts myofilament function and signaling cascades that target myofilaments (6–10). Although these studies were instrumental in identifying CapZ as a potential player in the acute response to exercise, intermittent stretching of muscle and the use of cultured cells fails to recapitulate the complex neural, humoral, immunological, and mechanical changes associated with aerobic exercise (2). Thus, despite the recognized importance of CapZ in cellular structure and function, its regulation under physiological conditions remains largely unexplored (11).

Several intracellular regulators of cardiac CapZ have been identified but their responses to physiological stressors like exercise have not been investigated. CapZ interacting protein (CapZIP) antagonizes the capping of filamentous actin by CapZ, an effect that is itself prevented when CapZIP is phosphorylated (4). By contrast the Hsc70-BAG3 complex stabilizes actin binding by CapZ (12). Eyers (4) showed that in skeletal muscle the CapZIP regulatory system is rapidly activated by electrical stimulation and allows for skeletal muscle actin remodelling. Whether this regulatory system is similarly affected in cardiac muscle is not known, nor has the impact of exercise on Hsc70-BAG3 been investigated.

It is unknown if myofilament remodelling takes place during the course of exercise, as previous studies have all examined changes post-exercise. Given the central role of CapZ in contributing to the architecture of the thin filaments, knowledge of its regulation is crucial for understanding the mechanisms by which the heart responds to exercise. The objectives of this study were to determine how sarcomeric CapZ is impacted by an acute bout of exercise and to identify the underlying molecular mechanisms that drive thin filament remodelling.

## Methods

### Animal Care and Use

Mice were group housed at the Central Animal Facility at the University of Guelph on a 12 h light/dark cycle and provided food and water ad libitum. All procedures were approved by and conducted in accordance with the guidelines set by the Animal Care and Use Committee of the University of Guelph and the Canadian Council on Animal Care.

### CapZ Deficient Mice

Cardiac-restricted CapZ-deficient mice have been described previously (7, 8, 13, 14). Z-disc-associated CapZ is decreased by ~20% through the overexpression of the β_2_-subunit specifically in the heart (13). This method reduces the amount of Z-disc associated with cardiac Z-disc (13). CapZ-deficient transgenic mice were homozygous for the transgene allele and were backcrossed to wildtype C57BL/6 mice for between 3 and 8 generations. All studies used female mice that were 8-12 weeks old. Wildtype mice were age, gender, and strain (C57BL/6) matched.

### Swimming Protocol

Mice were placed in a temperature controlled (30°C) water bath for 20 min and water recirculated to stimulate swimming. Immediately after exercise mice were euthanized; their hearts excised, rinsed in ice-cold saline (0.9% NaCl), snap frozen with liquid nitrogen, and stored at −80°C. For control mice water levels were decreased to allow mice to stand in water without swimming. Water temperature was maintained to ensure consistent temperature exposure between control and exercise groups.

### Myofilament Isolation

Cardiac myofilaments were isolated as we have published (9, 10). Briefly, hearts were homogenized in ice cold Standard Buffer (60 mM KCl, 30 mM Imidazole (pH 7.0), 2 mM MgCl_2_) containing phosphatase and protease inhibitors. The homogenate was centrifuged for 15 min at 12,000g at 4°C. The resulting pellet resuspended for 45 min in ice cold Standard Buffer plus 1% Triton X-100 and centrifuged at 1,100g for 15 min at 4°C. Pellets were washed three times in ice cold Standard Buffer. Protein concentration was measured with a Bradford Protein Assay (Bio-Rad Laboratories Ltd., Mississauga, ON).

### Immunoblotting

Isolated cardiac myofilaments were subjected to SDS-PAGE (12% separating gels) and transferred to nitrocellulose membranes for immunoblotting. Membranes were blocked in TBS containing 5% milk powder for 1 h at room temperature and then incubated with primary antibodies for CapZIP, phosphorylated CapZIP (S179, S108), BAG3, Hsc70, and CapZ overnight at 4°C. Horseradish peroxidase-conjugated secondary antibodies (anti-mouse, anti-rabbit, 1:5000) were added for 1 h at room temperature. Protein bands were detected using Western Lightning (PerkinElmer Life and Analytical Sciences, Woodbridge, ON) and a Bio-Rad ChemiDoc MP Imaging System (Bio-Rad Laboratories Ltd., Mississauga, ON). Data were analysed with ImageJ software (NIH, Bethesda, MD).

### Myofilament Protein Phosphorylation

Myofilament proteins (10 μg) were separated using 12% SDS-PAGE and fixed in 50% methanol-10% acetic acid overnight at room temperature. Gels were stained with Pro-Q Diamond phosphoprotein stain (Thermo Fisher Scientific, Mississauga, ON) according to the manufacturer’s instructions to determine total protein phosphorylation. Gels were imaged using a Bio-Rad ChemiDoc MP Imaging System (Bio-Rad Laboratories Ltd., Mississauga, ON) and data analysed with ImageJ software (NIH, Bethesda, MD). Total protein load was determined by Coomassie staining of the same gels and total protein phosphorylation was normalized to total protein.

### Myofilament Activity

Actomyosin MgATPase activity was determined using a modified Carter assay as we have published (9). Isolated cardiac myofilaments (25 μg) were incubated in reaction buffers containing variable levels of free calcium. Activating (pCa 4.0) and relaxing (pCa 9.0) buffers were mixed to prepare the reaction buffers used. Myofilaments were incubated in reaction buffers for 10 min at 32°C and the reactions quenched with equal volumes of ice cold 10% trichloroacetic acid. The production of inorganic phosphate by ATP consumption was measured by adding 0.5% FeSO_4_ and 0.5% ammonium molybdate in 0.5 M H_2_SO_4_.

### Exercise Fatigue Testing

Exercise tolerance was determined by swimming or running mice to exhaustion. For the swim test mice were acclimated to the swimming bath with a 20 min swim exposure. Two days later mice were weighed and weights totalling 10% of body weight were attached to the tail. Mice were placed in a temperature controlled (30°C) water bath and water recirculated to stimulate swimming. Exhaustion was taken as the point when mice stopped swimming and sank to the bottom for ~5 s. For the running test mice were acclimated with two 10 min exercise sessions (separate 2 days) on the treadmill at a 5% grade incline, paced at 15 m/min. Fatigue testing began with a treadmill speed of 12 m/min on an incline of 40% and pacing was increased by 1 m/min at 2, 5, 10 and every subsequent 10 min (15). Exhaustion was defined as an inability/refusal of the mouse to continue running when gently encouraged with a bottle brush.

### Statistical Analysis

The effect of exercise on CapZ regulation, myofilament function, protein phosphorylation, and exercise performance were analysed using an unpaired Student’s t-test. All effects are compared to controls of the same animal group. Data are presented as mean ± SEM. P<0.05 was selected to indicate significant differences.

## Results

### Acute exercise impacts myofilament CapZ regulation

Cardiac myofilaments were isolated from mouse hearts following 20 min of swim exercise and CapZ levels measured by immunoblotting. Total myofilament CapZ protein levels were unaffected by exercise (Figure 1).

**Figure 1.**
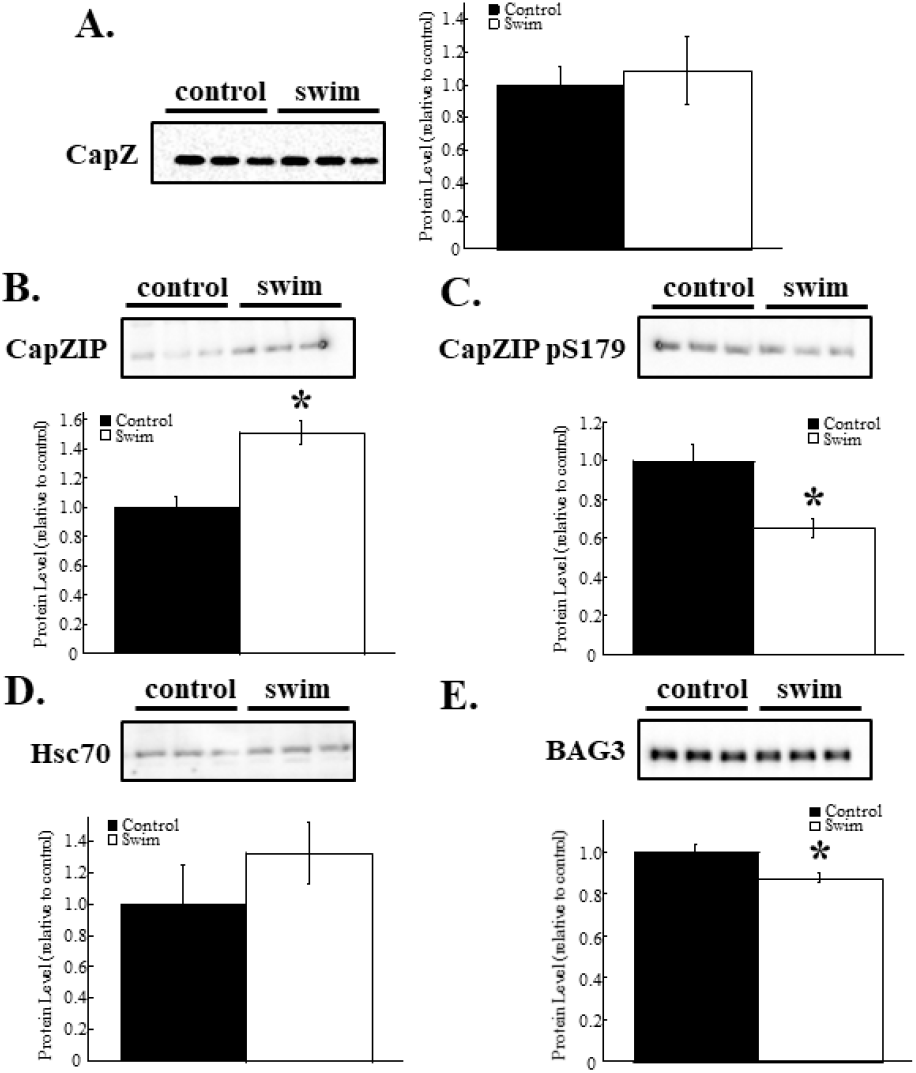
Cardiac CapZ regulation following acute exercise. Cardiac myofilaments were isolated after an acute bout of exercise and probed with immunoblotting. **A.** Myofilament CapZ levels were not impacted by an acute bout of swimming. **B.** CapZIP increased 51 ± 8% over control samples. **C.** Phosphorylation of CapZIP at S179 decreased by 35 ± 5% compared to non-exercise controls. **D.** Myofilament-associated Hsc70 levels were not altered by exercise. **E.** BAG3 decreased by 12 ± 2% compared to controls. N=9 in each group. Data are presented as mean ± SEM. *P<0.05 vs control.

CapZ can "wobble" when bound to actin by releasing the α-subunit from actin and remaining attached to the thin filament through the C-terminal tentacle of the β-subunit (16, 17). To determine if CapZ binding is affected by exercise, cardiac myofilaments from mice were probed with antibodies for the CapZ regulatory proteins CapZIP, Hsc70, and BAG3 following exercise. CapZIP protein increased 51 ± 8% over sedentary controls (Figure 1). Phosphorylation of myofilament-associated CapZIP at S179 was significantly decreased by 35 ± 5% with exercise. The increased levels of myofilament-associated CapZIP coupled with increased phosphorylation results in a weakened interaction between CapZ and actin (4). Hsc70 and BAG3 strengthen CapZ-actin interactions (12). Myofilament Hsc70 levels increased by 32 ± 20% with exercise, although this was not significantly different than controls (p=0.32). The Hsc70 binding partner BAG3 decreased 12 ± 2% with exercise. The lack of any coherent changes in Hsc70-BAG3 levels indicates that this complex is unlikely to affect CapZ binding to sarcomeric actin.

### Reduced cardiac CapZ shortens time to fatigue

To determine what role CapZ has in the acute response to exercise we subjected mice that have an approximately 20% reduction in cardiac CapZ levels (7, 8) to exhaustive bouts of exercise. In a swimming protocol the time to exhaustion for wildtype mice was 682 ± 78 s (Figure 2A). CapZ-deficient transgenic mice had a significantly shorter time to exhaustion with an average of 454 ± 65 s. Similarly, in a running protocol the time to exhaustion was 5036 ± 396 s for wildtype mice, which was significantly longer than the 4531 ± 178 s of cardiac CapZ-deficient transgenic mice (Figure 2B). The results demonstrate that a loss of myofilament-associated CapZ impairs exercise performance.

**Figure 2.**
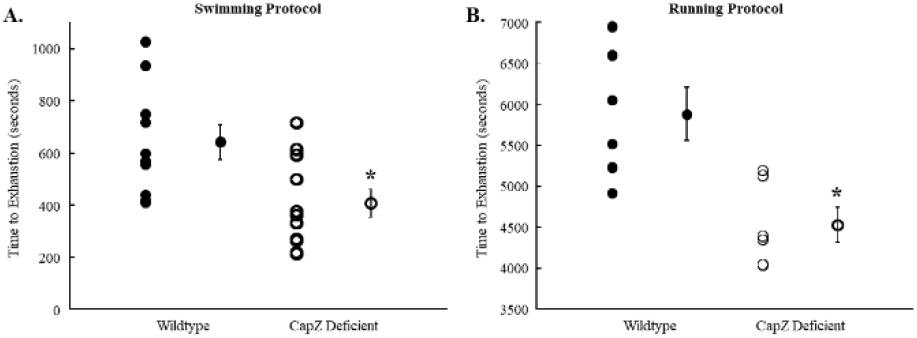
Time to exhaustion in wildtype and cardiac CapZ- deficient mice. Mice were exercised to exhaustion using a **A.** Swimming or **B**. Running test. Average time to exhaustion for wildtype mice using the swimming protocol was 682 ± 78 s which was significantly longer than the 454 ± 65 s for CapZ-deficient transgenic mice. The running protocol produced similar results with wildtype mice taking 5036 ± 396 s to reach exhaustion, compared to only 4531 ± 178 s for cardiac CapZ-deficient transgenic mice. N=10 in each group for swimming and N=6 in each group for running. Data are presented as mean ± SEM. *P<0.05 vs control of the same exercise.

### Reduced CapZ impacts CapZ regulation

To determine if CapZ regulation during exercise was disrupted in cardiac CapZ-deficient mice, cardiac myofilaments were probed for CapZ and key regulatory proteins using immunoblotting. CapZ, CapZIP, and Hsc70 levels were unchanged following exercise, as was the phosphorylation of CapZIP (Figure 3). Myofilament BAG3 levels increased 10 ± 3% following exercise. Unlike wildtype mice, CapZ-deficient transgenic mice lack a CapZ regulatory response to acute exercise, as measured by changes in myofilament CapZIP or Hsc70-BAG3.

**Figure 3.**
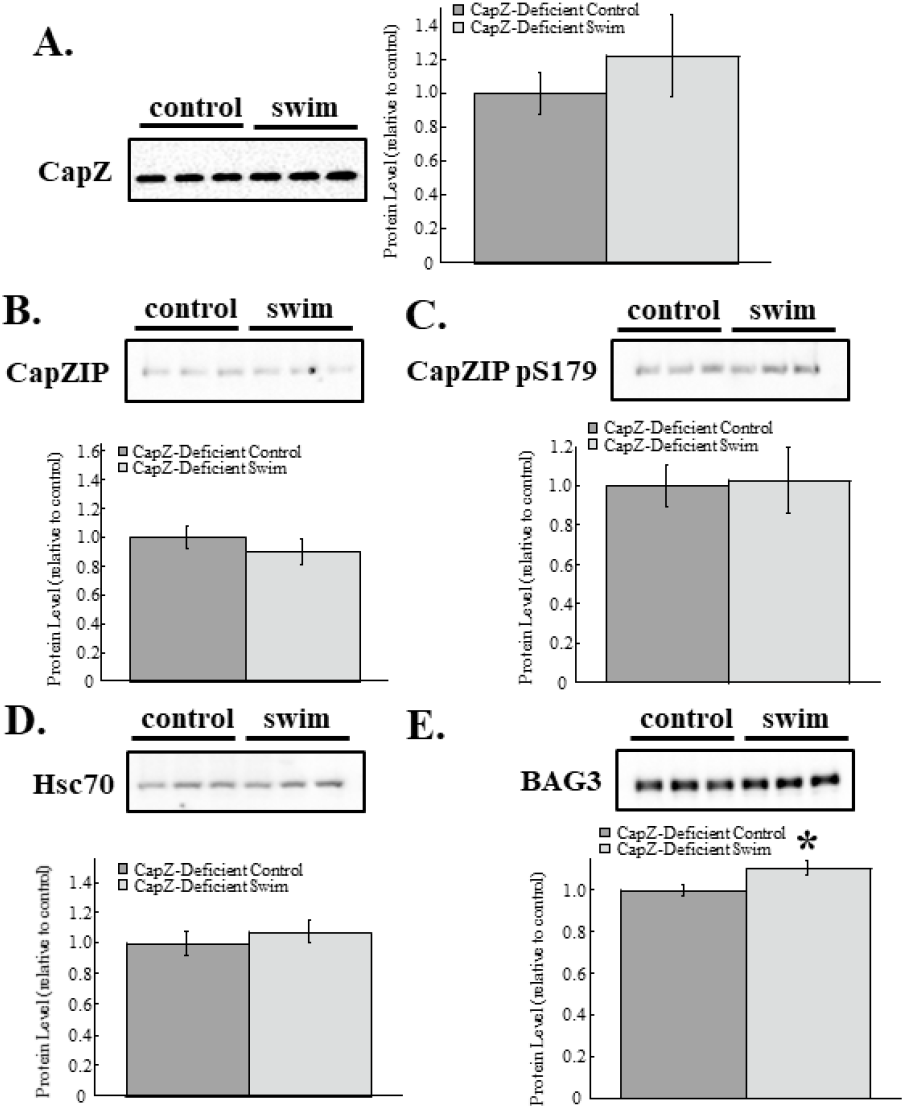
Cardiac CapZ-deficient mice exhibit an impaired CapZ regulatory response to exercise. Cardiac myofilaments were isolated after an acute bout of exercise and probed with immunoblotting. Neither **A.** CapZ, **B**. CapZIP, **C.** phosphorylation of CapZIP at S179, nor D. myofilament-associated Hsc70 levels were affected by exercise. **E**. BAG3 increased by 10 ± 3% compared to controls. N=9 in each group. Data are presented as mean ± SEM. *P<0.05 vs control.

### Reduced CapZ alters myofilament functional response to exercise

Cardiac myofilaments are the central contractile elements of the heart. Myofilament function was measured using actomyosin MgATPase activity following an acute bout of submaximal swimming. Myofilament function in wildtype hearts was not significantly affected by exercise (Figure 4A). Cardiac myofilaments from CapZ-deficient mice revealed a significant reduction in function at saturating levels of calcium and at several points of submaximal activation (<1 μM free calcium) (Figure 4B). These data indicate that reduced CapZ impairs the ability of cardiac myofilaments to maintain function in the face of an exercise stressor.

**Figure 4.**
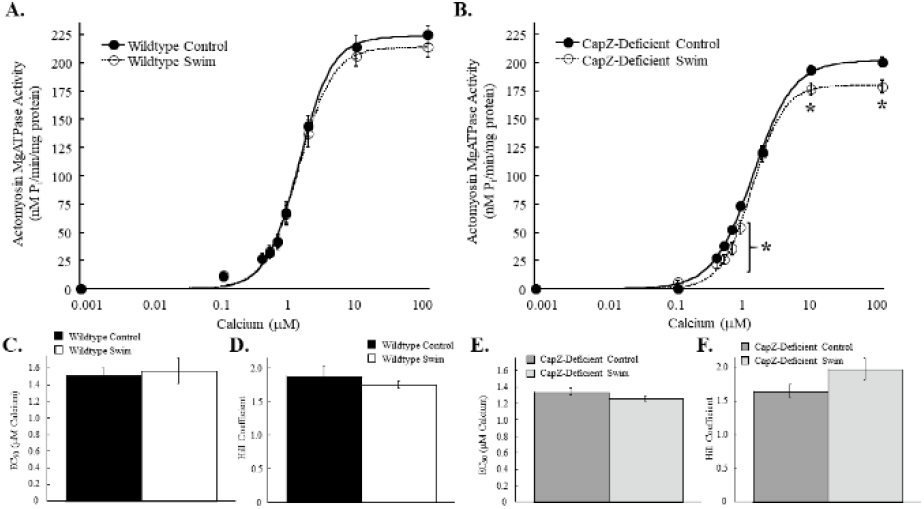
Myofilament actomyosin MgATPase activity with a single bout of exercise. Wildtype and cardiac CapZ-deficient transgenic mice were subjected to a single bout of exercise and myofilament function assessed with an actomyosin MgATPase assay. **A.** Actomyosin MgATPase-calcium relationship was not altered with exercise in wildtype myofilaments. **B**. Exercise decreased maximum actomyosin MgATPase activity in cardiac CapZ-deficient transgenic mice from 223 ± 5 nM P/min/mg protein to 199 ± 6 nM P/min/mg protein. At free calcium concentrations below 1 μM actomyosin MgATPase activity was significantly impaired in cardiac CapZ-deficient myofilaments. In wildtype myofilaments neither **C.** EC_50_ nor **D**. Hill coefficient were significantly impacted by exercise. Similarly, both **E**. EC_50_ and **F**. Hill coefficient were not significantly altered by exercise in cardiac CapZ- deficient myofilaments. N=9 in each group. Data are presented as mean ± SEM. *P<0.05 vs control for same group. **Key:** MyBP-C, myosin binding protein C; MLC2, myosin light chain 2; Phosphorylation, ProQ Diamond stained gels for total phosphorylation; Coomassie, Coomassie stained gels for total protein load.

Phosphorylation of cardiac myofilaments permits a rapid response to sudden onset stressors like exercise. Cardiac myofilaments were resolved by SDS-PAGE and phosphorylation levels probed with ProQ Diamond staining following a submaximal swim ming protocol. Myofilaments from wildtype mice exhibited significant increases in total phosphorylation levels for myosin binding protein C, desmin, troponin T, tropomyosin, and troponin I, whereas myosin light chain 2 was not significantly affected (Figure 5A and 5C). CapZ-deficient myofilaments showed no significant changes in total phosphorylation although troponin I (p=0.11) and myosin light chain 2 (p=0.10) phosphorylation levels tended to rise (Figure 5B and 5D).

**Figure 5.**
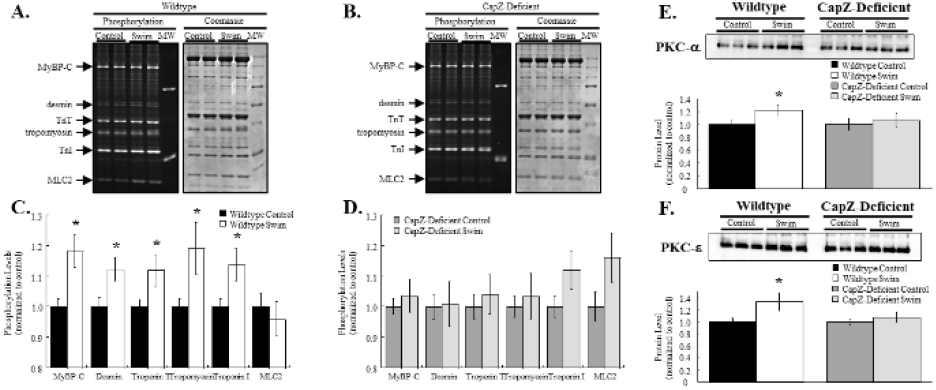
Changes in myofilament protein phosphorylation following a single bout of exercise. Following exercise cardiac myofilaments were resolved by SDS-PAGE and stained for total phosphorylation. **A.** Myofilaments from wildtype mice had significant increases in MyBP- C, desmin, troponin T, tropomyosin, and troponin I, while MLC2 was unaltered. **B.** By contrast there were no significant phosphorylation changes found in cardiac myofilaments from CapZ-deficient transgenic mice. Summary results are shown in panels **C** and **D**. Myofilament levels of PKC-α increased after exercise in wildtype mice but not CapZ-deficient transgenic mice. **E**. Similarly, myofilament levels of PKC-ε also increased in wildtype mice post-exercise but not in CapZ-deficient transgenic mice. Representative images of phosphorylation gels, Coomassie stained gels, and immunoblots are shown. N=9 in each group. Data are presented as mean ± SEM. *P<0.05 vs control for same group.

Cardiac myofilament phosphorylation is mediated by a number of different kinases. Work by us (7–10, 14) and others (18–21) have found that CapZ influences myofilament regulation by protein kinase C (PKC) and is a substrate for this family of kinases. To determine how exercise alters myofilament-associated PKC and if a decrease in cardiac CapZ levels alters exercise-dependent changes in sarcomeric PKC we probed isolated cardiac myofilaments for PKC-α (calcium-dependent isoform) and PKC-ε (calcium independent isoform). Whereas acute exercise increased both myofilament-associated PKC-α and -ε in wildtype hearts, the transgenic reduction in CapZ levels prevented these increases (Figure 5E and 5F). These patterns of PKC activation may explain both the increase in myofilament protein phosphorylation following exercise in wildtype mice and the lack of change in CapZ-deficient mice.

### Reduced CapZ destabilizes sarcomeric actin filaments during acute exercise

Exercise imposes a physical and neurohumoral stress on cardiac myocytes and demands a higher level of heart function. CapZ regulates both actin filament growth and acts as a physical anchor for the thin filaments at Z-discs. To determine if the reduction in CapZ destabilizes the myofilaments under these stressors we probed isolated cardiac myofilaments for anchoring and stabilizing proteins after an acute bout of swimming. Total sarcomeric actin was unaffected by submaximal exercise in both wildtype and CapZ-deficient transgenic mice (Figure 6). The pointed-end actin binding protein tropomodulin was also unaffected by exercise in both groups. Levels of the titin capping protein telethonin/Tcap did not change in wildtype mice, but significantly decreased by 39 ± 8% in CapZ-deficient mice. The actin-binding protein leiomodin 2 increased 78 ± 28% in myofilaments from wildtype mice following acute exercise whereas myofilaments from CapZ-deficient mice exhibited no significant changes in leiomodin 2 levels. Together these results show that cardiac myofilament regulation by a variety of binding proteins is disrupted with a small decrease in CapZ levels.

**Figure 6.**
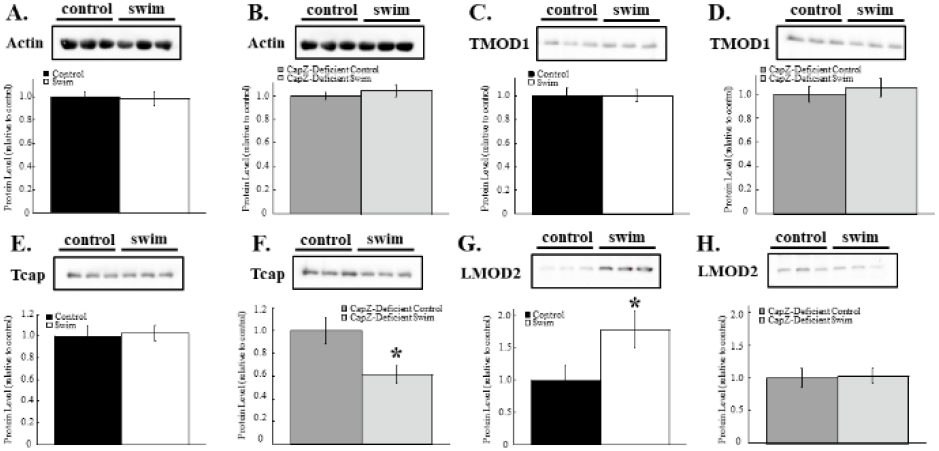
Myofilament-binding proteins following acute exercise. Cardiac myofilaments were isolated after a single exercise session and subjected to immunoblotting. Sarcomeric actin levels were not altered by exercise in wildtype or cardiac CapZ-deficient transgenic mice (**A** and **B**). Similarly, tropomodulin levels remained unaffected by exercise (**C** and **D**). Tcap/telethonin levels did not change with exercise in wildtype myofilaments **(E)** but did decrease by 39 ± 8% in CapZ-deficient mice **(F)**. Conversely, leiomodin levels rose 78 ± 28% in myofilaments from wildtype mice with exercise **(G)** but did not change in cardiac CapZ-deficient mice **(H)**. N=9 in each group. Data are presented as mean ± SEM. *P<0.05 vs control for same group. **Key:** TMOD1, tropomodulin 1; Tcap, telethonin; LMOD2, leiomodin 2.

## Discussion

Exercise exerts a rapid stress on the heart that must be quickly matched to maintain performance and prevent cardiovascular collapse. The molecular bas is of these changes are poorly characterized, in particular myofilament adaptations, limiting our understanding of how the heart responds to physiological stress. In this study we identify the actin capping protein CapZ as a critical player in the cardiac response to exercise. CapZ binding to sarcomeric actin is weakened during exercise, and yet its reduction limits exercise capacity in mice, indicating a finely controlled remodelling of the thin filaments. The presence of CapZ during exercise is necessary to maintain normal myofilament function in the face of increased stress as demonstrated by a decrease in exercise performance when CapZ expression is reduced. Finally, a decrease in CapZ expression disrupts the binding of several critical myofilament-associated proteins during exercise like Tcap/telethonin and leiomodin, which help to maintain sarcomeric integrity. Together this study represents the first investigation of molecular changes in cardiac myofilaments during an acute bout of exercise and discovers a critical role for CapZ in the myofilament adaptation to exercise.

CapZ exists in a ‘wobble’ state with actin in which the α-subunit of the capping protein dynamically binds and dissociates from actin, remaining bound only through the β-subunit (16). In situations like cell motility where actin uncapping periods may be prolonged to allow for sustained filamentous growth the complete dissociation of CapZ can occur. But in striated muscle where prolonged periods of unrestrained actin addition could interfere with contractile function this model is not likely. An analogous situation has been described for the pointed-end capping protein tropomodulin in which actin depolymerization is permitted without the dissociation of tropomodulin from the thin filament (22). Our data show that during exercise CapZ does not leave the myofilament compartment. However, the presence of increased levels of CapZIP would favour a disruption of the CapZ-actin binding. Based on these data we propose a model of CapZ regulation during exercise in which actin capping by CapZ is weakened by CapZIP. This weakening comes in the form of favouring the open wobble state where CapZ remains bound to actin through the β-subunit, but the uncapping may be sufficient to allow for actin additions. We found no change in the levels of myofilament actin during exercise which suggests that actin filament lengths are maintained during exercise, and that the weakening of CapZ allows for increased treadmilling of actin. Lin and colleagues (20) showed that mechanical perturbation of neonatal cardiac myocytes meant to mimic exercise induces a rapid increase in actin dynamics, which provides support for our model.

The impact of exercise training on the heart is well known. By contrast the molecular alterations that drive the myocardial response to acute exercise have not been widely investigated, in particular the role of cardiac myofilaments. Muller et al (23) found that a single bout of acute exercise increased passive stiffness of isolated cardiac myocytes through a complex alteration in titin phosphorylation. Although an increase in myofilament stiffness can contribute to diastolic dysfunction the acute changes in stiffness that occur in exercise were hypothesized to improve cardiac output by supporting the Frank-Starling mechanism of increased contractility (23). Chakouri and colleagues (24) examined post-translational changes in cardiac myofilaments subjected to exercise and found that a submaximal stress increased troponin I and myosin binding protein C phosphorylation at protein kinase A-preferred sites. Interestingly, these myofilament alterations did not impact myofilament activation as assessed by force-calcium curves. In our study we found a similar increase in myofilament protein phosphorylation and extend these changes to include desmin, troponin T, and tropomyosin, but not myosin light chain 2. We also found no significant changes in myofilament activation as quantified by an actomyosin MgATPase assay. Although there were no changes in myofilament function detectable in association with acute exercise this does not preclude an important role for cardiac myofilaments. Like the titin phosphorylation changes reported by Muller and colleagues, the increase in desmin phosphorylation we observed may provide physical support for the stress of exercise. Furthermore, the oxidative stress of exercise has negative effects on cardiac myofilaments and these changes may be countered by phosphorylation (24). In this scenario the increase in phosphorylation may offset the detrimental effects of myofilament oxidation, helping to maintain contractility.

The decrease in time to exhaustion of CapZ-deficient mice was associated with a significant decline in myofilament activation at several submaximal calcium points and at saturating levels. The change in maximum activation is unlikely to impact physiological function as intracellular calcium levels do not rise to this level (>100 μM) under normal circumstances, but the measurable change is a sentinel of myofilament targeting. The reduction in activation at submaximal calcium levels may explain the poorer exercise performances in CapZ-deficient mice. The relative reduction in myofilament calcium sensitivity at low levels of calcium exposure could slow myofilament activation. As heart rates rise in exercise these small changes in myofilament kinetics may significantly impair the ability of the myofilaments to contract in the relatively short activation win dow.

Acute exercise applies a sudden stress to the heart, necessitating a rapid response. Myofilament protein phosphorylation allows for a quick and powerful response to stressors. These changes are mediated by protein kinases and phosphatases that have a number of potential targets within the contractile complex. Muller has reported PKC-dependent increases in titin phosphorylation, which they ascribed to increased PKC-α activity, although they did not determine if PKC-α binding to cardiac myofilaments was altered with exercise. We have previously shown that CapZ is critical for PKC-dependent control of cardiac myofilaments and studies from Russell’s group showed that CapZ binding is itself regulated by PKC-ε (21, 25). In this study we show for the first time that myofilament-associated PKC-α and -ε increase during submaximal exercise and that these changes are mitigated in mice with a deficiency of cardiac CapZ. The inability of CapZ-deficient transgenic mice to maintain physical activity for as long as wildtype may, in part, be explained by the disruption in PKC signaling. Without PKC-α myofilament stiffness driven by titin phosphorylation would be reduced and the Frank-Starling effect impaired, while a lack of increased PKC-ε would impair CapZ regulation and thin filament remodelling.

Our data show that an acute bout of exercise is associated with a significant increase in cardiac myofilament-associated leiomodin in wildtype mice, but not CapZ-deficient transgenic animals. Leiomodin is a known actin nucleator that co-localizes with tropomodulin at the pointed-ends of actin filaments. Our data show no significant increase in sarcomeric actin which makes actin filament elongation unlikely, but a model of actin treadmilling proposed by Skwarek-Maruszewska and colleagues (26) may explain enhanced myofilament leiomodin in the absence of increased sarcomeric actin. In this model increased contractility stimulates leiomodin to bind to actin filaments near the Z-disc and remain bound as actin treadmills through the filament. Leiomodin binding stabilizes the thin filament structure and maintains contractility in the face of an increase in demand (26, 27). The ability of leiomodin to stabilize the dynamically changing myofilaments in exercising wildtype mice but not CapZ-deficient transgenic mice could explain the enhanced performance of wildtype mice over their CapZ-deficient counterparts.

Cardiac Z-discs, long considered to be passive structural elements of the sarcomere, have emerged as dynamic players in the contractile apparatus of the heart. In particular, we show here and in previous studies (6–10, 14) that the Z-disc protein CapZ is a critical regulator of myocardial function and intracellular signaling. Understanding the role of CapZ in shaping the myocardial response to physiological and pathological stress is critical to differentiating between the adaptive and maladaptive mechanisms that characterize exercise training and disease respectively. Insight into the mechanistic underpinnings of exercise offers a unique opportunity to target these pathways therapeutically to produce beneficial effects in the heart and to potentially mitigate disease.

## References

1. Vatner SF, Pagani M. Cardiovascular adjustments to exercise: hemodynamics and mechanisms. Prog Cardiovasc Dis. [date unknown];19(2):91–108.

2. Booth FW, Thomason DB. Molecular and cellular adaptation of muscle in response to exercise: perspectives of various models. Physiol Rev. 1991;71(2):541–85.

3. Kemi OJ, Ellingsen O, Smith GL, Wisloff U. Exercise-induced changes in calcium handling in left ventricular cardiomyocytes. Front Biosci. 2008;13(13):356.

4. Eyers CE, McNeill H, Knebel A, et al. The phosphorylation of CapZ-interacting protein (CapZIP) by stress-activated protein kinases triggers its dissociation from CapZ. Biochem J. 2005;389(Pt 1):127–35.

5. Li J, Russell B. Phosphatidylinositol 4,5-bisphosphate regulates CapZβ1 and actin dynamics in response to mechanical strain. Am J Physiol Heart Circ Physiol. 2013;305(11):H1614–23.

6. Gaikis L, Stewart D, Johnson R, Pyle WG. Identifying a role of the actin capping protein CapZ in β-adrenergic receptor signalling. Acta Physiol (Oxf). 2013;207(1):173–82.

7. Pyle WG, Hart MC, Cooper JA, Sumandea MP, de Tombe PP, Solaro RJ. Actin capping protein: an essential element in protein kinase signaling to the myofilaments. Circ Res. 2002;90(12):1299–306.

8. Pyle WG, La Rotta G, de Tombe PP, Sumandea MP, Solaro RJ. Control of cardiac myofilament activation and PKC-betaII signaling through the actin capping protein, CapZ. J Mol Cell Cardiol. 2006;41(3):537–43.

9. Yang FH, Pyle WG. Cardiac actin capping protein reduction and protein kinase C inhibition maintain myofilament function during cardioplegic arrest. Cell Physiol Biochem. 2011;27(3-4):263–72.

10. Yang F, Aiello DL, Pyle WG. Cardiac myofilament regulation by protein phosphatase type 1α and CapZ. Biochem Cell Biol. 2008;86(1):70–8.

11. Hernandez-Valladares M, Kim T, Kannan B, et al. Structural characterization of a capping protein interaction motif defines a family of actin filament regulators. Nat Struct Mol Biol. 2010;17(4):497–503.

12. Hishiya A, Kitazawa T, Takayama S. BAG3 and Hsc70 interact with actin capping protein CapZ to maintain myofibrillar integrity under mechanical stress. Circ Res. 2010;107(10):1220–31.

13. Hart MC, Cooper JA. Vertebrate Isoforms of Actin Capping Protein. J Cell Biol. 1999;147(6):1–12.

14. Yang FH, Pyle WG. Reduced cardiac CapZ protein protects hearts against acute ischemia-reperfusion injury and enhances preconditioning. J Mol Cell Cardiol. 2012;52(3):761–72.

15. Castellani LN, Peppler WT, Miotto PM, Bush N, Wright DC. Exercise Protects Against Olanzapine-Induced Hyperglycemia in Male C57BL/6J Mice [Internet]. Sci Rep. 2018;8(1) doi:10.1038/s41598-018-19260-x.

16. Bhattacharya N, Ghosh S, Sept D, Cooper JA. Binding of Myotrophin/V-1 to Actin-capping Protein. J Biol Chem. 2006;281(41):31021–30.

17. Kim K, McCully ME, Bhattacharya N, Butler B, Sept D, Cooper JA. Structure/function analysis of the interaction of phosphatidylinositol 4,5-bisphosphate with actin-capping protein: Implications for how capping protein binds the actin filament. J Biol Chem. 2007;282(8):5871–9.

18. Lin Y-H, Warren CM, Li J, McKinsey TA, Russell B. Myofibril growth during cardiac hypertrophy is regulated through dual phosphorylation and acetylation of the actin capping protein CapZ. Cell Signal. 2016;28(8):1015–24.

19. Mkrtschjan MA, Solís C, Wondmagegn AY, Majithia J, Russell B. PKC epsilon signaling effect on actin assembly is diminished in cardiomyocytes when challenged to additional work in a stiff microenvironment. Cytoskeleton (Hoboken). 2018;75(8):363–71.

20. Lin YH, Li J, Swanson ER, Russell B. CapZ and actin capping dynamics increase in myocytes after a bout of exercise and abates in hours after stimulation ends. J Appl Physiol. 2013;114(11):1603–9.

21. Hartman TJ, Martin JL, Solaro RJ, Samarel AM, Russell B. CapZ dynamics are altered by endothelin-1 and phenylephrine via PIP2- and PKC-dependent mechanisms. Am J Physiol Cell Physiol. 2009;296(5):C1034–9.

22. Gregorio CC, Fowler VM. Mechanisms of thin filament assembly in embryonic chick cardiac myocytes: Tropomodulin requires tropomyosin for assembly. J Cell Biol. 1995;129(3):683–95.

23. Muller AE, Kreiner M, Kotter S, et al. Acute exercise modifies titin phosphorylation and increases cardiac myofilament stiffness. Front Physiol. 2014;5:449.

24. Chakouri N, Reboul C, Boulghobra D, et al. Stress-induced protein S-glutathionylation and phosphorylation crosstalk in cardiac sarcomeric proteins - Impact on heart function. Int J Cardiol. 2018;258:207–16.

25. Lin YH, Swanson ER, Li J, Mkrtschjan MA, Russell B. Cyclic mechanical strain of myocytes modifies CapZβ1 post translationally via PKCε. J Muscle Res Cell Motil. 2015;36(4-5):329–37.

26. Skwarek-Maruszewska A, Boczkowska M, Zajac AL, et al. Different localizations and cellular behaviors of leiomodin and tropomodulin in mature cardiomyocyte sarcomeres. Mol Biol Cell. 2010;21(19):3352–61.

27. Szatmári D, Bugyi B, Ujfalusi Z, Grama L, Dudás R, Nyitrai M. Cardiac leiomodin2 binds to the sides of actin filaments and regulates the ATPase activity of myosin. PLoS One. 2017;12(10):e0186288.

